# Synthesis of C8-vinyl chlorophylls *d* and *f* impairs far-red light photoacclimation and growth under far-red light

**DOI:** 10.1101/2025.05.26.656186

**Authors:** Afeefa K. Chaudhary, Lorenz K. Fuchs, Krzysztof Pawlak, Gregory F. Dykes, Royston Goodacre, Dennis J. Nürnberg, Daniel P Canniffe

## Abstract

The inducible biosynthesis of chlorophylls *d* and *f* enables a subset of specialised cyanobacteria to perform oxygenic photosynthesis under far-red light—in the absence of visible wavelengths—via a process termed far-red light photoacclimation. These pigments, like the more common chlorophylls *a* and *b*, typically carry an ethyl substituent at the C8 position of the macrocycle, formed by reduction of a vinyl group by an 8-vinyl reductase enzyme. Here, we disrupted the gene encoding BciB, an 8-vinyl reductase found in the majority of cyanobacteria, in *Chroococcidiopsis thermalis* PCC 7203, a model for the study of far-red light photoacclimation. Disruption of *bciB* results in the synthesis of 8-vinyl chlorophyll *a* when the cells are grown in white light; on switching to far-red light, synthesis of 8V-forms of chlorophylls *d* and *f*, which have not been detected in nature, are synthesised in this strain. The *bciB* mutant exhibits sensitivity to high irradiance under both light regimes. Pigment analysis and whole-cell absorption and fluorescence spectroscopy reveal decreased synthesis of far-red absorbing chlorophylls, reduced photosystem assembly and an impaired acclimation to far-red light, and transmission electron microscopy demonstrates altered thylakoid membrane morphology in the mutant when compared to the wild type. These results demonstrate the importance of 8-vinyl group reduction for acclimation to far-red light.

## INTRODUCTION

Photosynthesis, the process by which sunlight is captured and converted into the chemical energy that supports life on Earth, is dependent on the chlorophyll (Chl) family of pigments. These Chls act as antenna chromophores to capture photons and transfer derived excitation energy towards the photochemical reaction centre, where they are also involved in charge separation and the primary steps of photosynthetic electron transport [1]. Chl *a* is the ubiquitous pigment essential for oxygenic photosynthesis and is the sole Chl found in the majority of cyanobacteria. In addition to Chl *a*, plants, algae, and a few cyanobacteria in the Prochlorococcaceae, Prochlorotrichaceae (together formerly known as Prochlorales or prochlorophyta) and a single member of the Acaryochloridaceae synthesize Chl *b* [2,3], which carries a formyl group at C7 (**Fig. 1**), enhancing light capture in the blue and orange regions of the solar spectrum [4]. In 1993, a cyanobacterium was isolated from a colonial sea squirt off the coast of Palau that was found to contain Chl *d*, carrying a formyl group at C3 (see **Fig. 1**), as its major photopigment [5]. Chl *d* has a red-shifted absorption maximum compared to that of Chl *a*, displaying a Q_y_ band at 696 nm—31 nm longer than that of Chl *a*—and the high Chl *d*:Chl *a* ratio *in vivo* (>90% Chl *d*) accounts for the strongly red shifted absorption profile of this organism, *Acaryochloris marina*, relative to the majority of cyanobacteria, allowing it to capture light at wavelengths > 700 nm in the solar spectrum (far-red light; FRL) [5]. In 2010, another novel pigment, Chl *f*, carrying a formyl group at C2 resulting in a Q_y_ absorption maximum at 705 nm (see **Fig. 1**), was identified in a stromatolite from Shark Bay, Australia [6]. This pigment was accumulated by a cyanobacterium, *Halomicronema hongdechloris*, enriched by growth under FRL—conditions that mirror its ecological niche in the stromatolite—but Chl *f* in this cyanobacterium was found to be a minor component, making up 10% of the total Chl when grown under FRL, and undetectable in white light (400–700 nm; WL) [6,7]. Subsequently, this induction of the synthesis of Chl *f* upon growth in FRL was discovered in another cyanobacterium, *Leptolyngbya* sp. JSC-1, isolated from a floating microbial mat at 45°C close to La Duke Hot Spring, Montana, USA [8]. In this case it was discovered that Chl *f* synthesis occurred concomitant with synthesis of Chl *d* and the replacement of the core subunits of photosystems I (PSI), II (PSII), and the phycobilisome antenna with paralogous proteins encoded in a gene cluster, expression of which is controlled by a red/far-red phytochrome and two response regulators [8]. This response to growth in FRL, termed ‘far-red light photoacclimation’ (FaRLiP), results in the increase in expression by more than two times for ∼900 genes, and a decrease to less than half for ∼2000 genes, accounting for a two-fold change in expression of >40% of the genome [8]. Soon after, the ability to perform FaRLiP was discovered in morphologically diverse cyanobacteria, the majority from terrestrial environments in which competition for WL is fierce (e.g. microbial mats) or where FRL is enriched due to scattering of sunlight (e.g. soils) [9]; In these diverse organisms the FaRLiP response is dependent on a conserved cluster of 19 core genes (although some clusters contain a greater number) [10]. Each FaRLiP cluster contains two paralogs of the PSII core subunit gene *psbA*, one of which encodes a light-dependent enzyme for the synthesis of Chl *f* [11,12].

**Figure 1.**
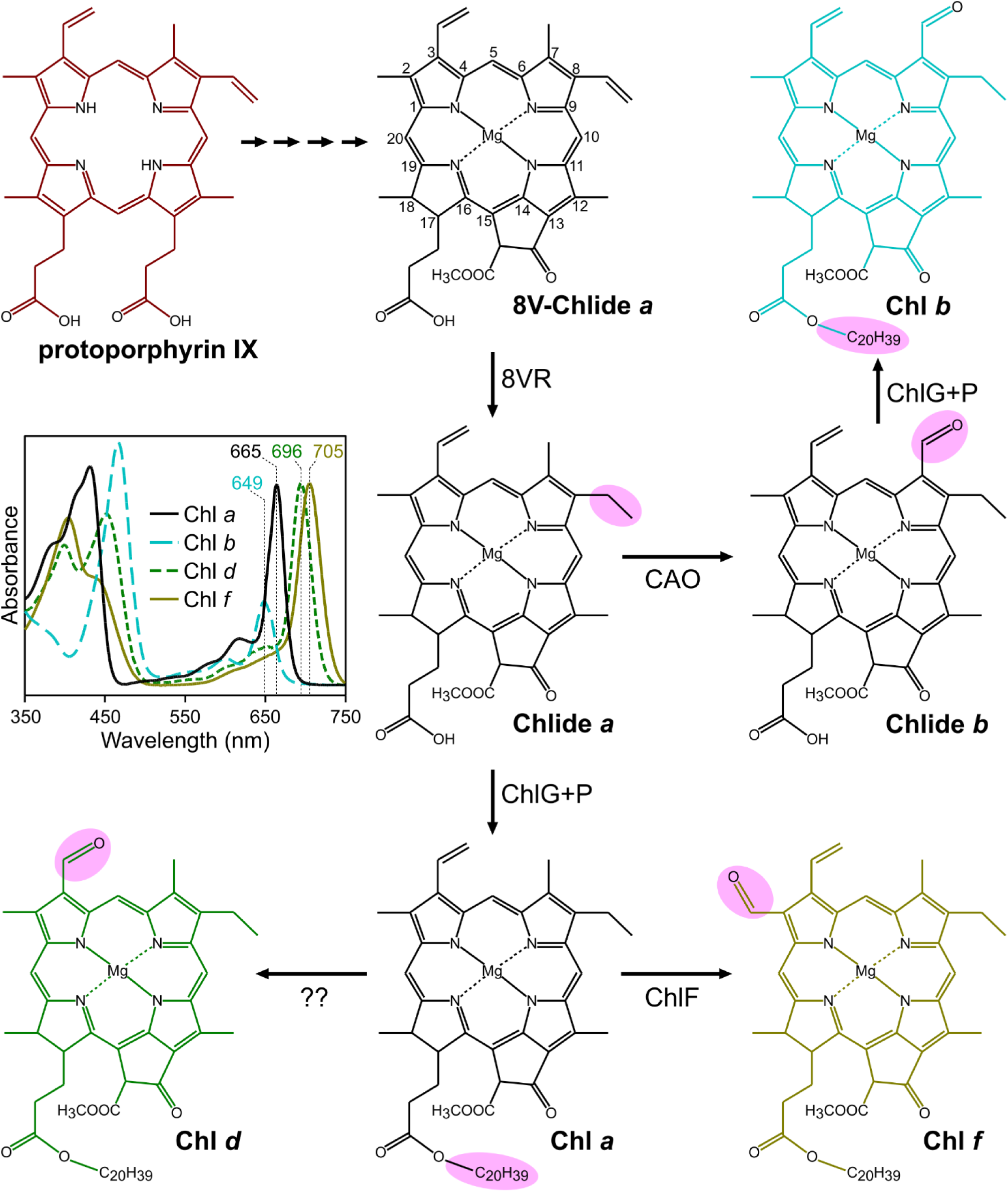
Terminal steps in the synthesis of Chls of oxygenic phototrophs. Protoporphyrin IX stands at the branchpoint between heme and Chl biosynthesis. IUPAC numbering of macrocycle carbons is displayed on 8V-Chlide, the universal precursor to all Chls. The routes to the synthesis of mature Chls are shown. The groups modified at each step are highlighted in pink, and the enzyme responsible is labelled at reaction arrows, except for Chl *d* synthase, which is currently unknown. CAO, chlorophyllide *a* oxygenase. 8VR, 8-vinyl reductase. ChlF, chlorophyll *f* synthase. ChlG, chlorophyll synthase. ChlP, geranylgeranyl reductase. Inset, absorption spectra of Chls are displayed, and their Q_y_ absorption maxima in methanol are labelled.

The mature pigments discussed above carry an ethyl group at the C8 position (8E), resulting from the reduction of a vinyl group on 8-vinyl (8V) chlorophyllide (Chlide) by an 8-vinyl reductase (8VR) enzyme (see **Fig. 1**); the only known exception to this are the marine *Prochlorococcus* spp. that naturally produce 8V-Chls *a* and *b* (also known as divinyl (DV)-Chl *a* and *b*) [13]. Outside of this special case, two unrelated classes of 8VR involved in the biosynthesis of chlorophylls are known to exist, BciA and BciB. BciA (EC 1.3.1.75) was first identified by screening mutants of *Arabidopsis thaliana*; mutations in the AT5G18660 locus led to the accumulation of 8V-rather than 8E-Chls *a* and *b* [14,15], and the recombinant enzyme produced in *Escherichia coli* was shown to reduce 8V-Chlide to 8E-Chlide [14]. Subsequently, BciA activities were demonstrated for proteins from rice [16], maize and cucumber [17], as well as in bacteriochlorophyll (BChl) *a*-synthesizing anoxygenic phototrophs including the green sulfur bacterium *Chlorobaculum tepidum* [18], and the purple phototrophic bacterium *Rhodobacter sphaeroides* [19]. *In vitro* assays performed with BciA-type 8VRs from various species have demonstrated that NADPH is the electron donor for this enzyme [16–18,20].

Despite also utilizing 8E-Chls, the genomes of most cyanobacteria do not contain orthologs of *bciA*, which indicated the existence of a second, unrelated 8VR. Two studies on the model cyanobacterium *Synechocystis* sp. PCC 6803 (*Synechocystis*) demonstrated that mutants in open reading frame (ORF) slr1923 were unable to grow under moderate light intensity and accumulated 8V-Chl *a* [21,22]. Subsequently, an ortholog of slr1923 from the green sulfur bacterium *Chloroherpeton thalassium* was shown to complement a *bciA* mutant of *Chlorobaculum tepidum*, recovering synthesis of 8E-(B)Chls and confirming the activity of the second class of 8VRs, annotated as BciB (EC 1.3.7.13) [23]. A study on the *in vitro* activity of the BciB-type 8VR from *Chloroherpeton thalassium* showed that the enzyme is a flavin adenine dinucleotide (FAD)-containing [4Fe–4S] protein, deriving electrons from reduced ferredoxin [24].

8VR mutants in plants and cyanobacteria display disrupted thylakoid membrane development and light sensitivity [14,16,21]. 8V-Chls *d* and *f* have never been reported in nature, so the effect that synthesis of these pigments have on the utilization of FRL is unknown. Species of *A. marina* appear to be unique among cyanobacteria by employing both BciA and BciB to ensure production of Chl *d* with an 8E group [25]. In the present study we constructed a *bciB* mutant of the model FRL-absorbing cyanobacterium *Chroococcidiopsis thermalis* PCC 7203 (hereafter *C. thermalis*) to assess the effect of 8V-Chls *d* and *f* on the FaRLiP process and growth under wavelengths longer than 700 nm.

## MATERIALS & METHODS

### Growth of bacterial strains

*C. thermalis* was grown photoautotrophically in BG-11 medium [26] buffered with 10 mM 2-[(2-Hydroxy-1,1-bis(hydroxymethyl)ethyl)amino] ethanesulfonic acid (TES) pH 8.2. Cells were routinely grown and maintained on solid medium containing 1.5% (w/v) Bacto agar or in liquid under moderate light intensity (30 μmol photons·m^−2^·s^−1^) provided by white-light LEDs at 30°C; when required, high light was provided by the same LEDs at 250 μmol photons·m^−2^·s^−1^. FRL was provided by LEDs with emission centred at 730 nm (Shenzhen Gouly LED Limited); moderate and high intensity FRL was calculated at equivalents of 20 and 75 μmol photons·m^−2^·s^−1^, respectively. Light intensity was measured with a LI-COR LI-250 radiometer equipped with a LI-190R pyranometer visible–near infrared sensor. When necessary, BG-11 medium was supplemented with 20 μg·ml^−1^ erythromycin.

*E. coli* strains were grown in lysogeny broth (LB) at 37°C in a rotary shaker at 250 rpm. When necessary, LB was supplemented with 30 μg·ml^−1^ spectinomycin, 100 μg·ml^−1^ ampicillin, and 12.5 μg·ml^−1^ chloramphenicol. All strains and plasmids used in this study are listed in Table S1.

### Plate-based growth assays

Drop growth assays of *C. thermalis* strains were conducted using liquid cultures grown under moderate light, adjusted to OD_780_ = 0.4 after Dounce homogenisation, and serially diluted. Dilutions were spotted onto solid medium and incubated at moderate and high (250 μmol photons·m^−2^·s^−1^) light intensity.

### Mutagenesis of *C. thermalis*

DNA sequences of ∼2.5 kb flanking the targeted sites for insertion of an antibiotic resistance cassette were amplified from the genome of *C. thermalis* with relevant UpF and UpR/DownF and Down R primer pairs (see Table S2 for all primer sequences) and fused by overlap extension PCR, such that the flanking regions were separated by a sequence containing XhoI and AvrII restriction sites. These fragments were digested with EagI and SacI and cloned into similarly digested pRL277 [27]. The erythromycin resistance cassette from pRL692 [28] was amplified with EmRF and EmRR, digested with XhoI and AvrII, and ligated into the same sites introduced by the overlap extension step in the constructed pRL277 plasmids. These plasmids were sequenced and transformed into the *E. coli* donor strain HB101, prepared to contain conjugal and helper plasmids pRL443 [29] and pRL528 [30], respectively, for biparental mating. Resulting HB101 transformants were grown overnight; 20 ml of culture was pelleted by centrifugation at 3000 x *g*, and the pellet was gently resuspended in 10 ml sterile LB. This was repeated three times to remove all traces of antibiotics, and the cells were finally resuspended in 200 µl LB. Meanwhile, 30 ml of freshly grown *C. thermalis* for each mating was sonicated for 2 x 30 s at low power to disperse clumps of cells and centrifuged as above. The pellet was washed with 10 ml sterile BG-11 medium three times and finally resuspended in 200 μl BG-11. These cells were mixed with the 200 μl suspension of HB101 cells in LB and incubated in the dark at RT for 8 h, before being spotted onto a sterile Whatman mixed cellulose ester filter membrane (0.45 μm pore size) overlaid on a BG-11 agar plate lacking antibiotics, and incubated at 30°C under dim light (∼2 μmol photons·m^−2^·s^−1^) for 24 h, after which time the membrane was transferred to a BG-11 plate containing erythromycin and exposed to moderate light. Green colonies appeared after ∼5 weeks; these were patched onto fresh selective plates for screening by colony PCR using the relevant CheckF and CheckR primers, which bind outside the regions of homology used for recombination, and positive patches were serially streaked onto selective plates to isolate axenic colonies away from any remaining HB101 cells.

### Absorption spectroscopy

UV/vis/near-IR absorption spectra of homogenised whole cells were collected on a Cary 3500 spectrophotometer (Agilent Technologies) scanning between 300 and 1100 nm at 1 nm intervals with a 0.1 s integration time.

### Fluorescence spectroscopy

Cells were harvested from liquid medium and resuspended in BG-11 containing 60% (v/v) glycerol by gentle homogenisation. Cryogenic fluorescence emission spectra were measured at 77 K in an OptistatDN-X cryostat (Oxford Instruments) on an Edinburgh Instruments FLS1000 spectrofluorometer between 600 and 850 nm at 1 nm intervals and a 0.2 s dwell time, after excitation at 440 nm.

### Extraction and analysis of pigments

Chls were extracted from cell pellets after washing in 20 mM HEPES pH 7.2 by adding 9 pellet volumes of 7:2 (v/v) acetone: methanol, vortex mixing for 30 s, and incubating on ice for 15 min. Extracts were clarified by centrifugation (15,000 × *g* for 5 min at 4°C), and the supernatants were filtered through 0.22 µm PVDF membrane filters and immediately analysed by high-performance liquid chromatography (HPLC) on an Agilent 1100 system. Chl species were separated on a Supelco Discovery HS C18 column (5 µm particle size, 120 Å pore size, 250 × 4.6 mm) using a program modified from that of a previously published method [31]. Solvents A and B were 80:20 (v/v) methanol:500 mM ammonium acetate and 80:20 (v/v) methanol:acetone, respectively. Pigments were eluted at 1 ml·min^−1^ at 40°C on a linear gradient of 92 to 94% solvent B over 25 min, increasing to 100% to wash the column. Elution of Chl species was monitored by checking absorbance at 665 nm (Chl *a*), 696 nm (Chl *d*), and 706 nm (Chl *f*).

### Transmission electron microscopy

*C. thermalis* cells were prefixed with 2.5% (w/v) glutaraldehyde and 4% (w/v) paraformaldehyde in 0.1 M sodium cacodylate pH 7.2 and postfixed with 1% (w/v) osmium tetroxide, 1.5% (w/v) potassium ferrocyanide, 0.1% (w/v) thiocarbohydrazide, 2% (w/v) osmium tetroxide and 1% (w/v) uranyl acetate. Samples were dehydrated with increasing ethanol concentrations (30−100%), embedded in TAAB 812 premix resin, and cut into thin sections (70 nm). These were post-stained with 3% (w/v) lead citrate. Images were recorded using a Tecnai G2 Spirit BioTWIN transmission electron microscope (FEI) equipped with a Gatan Rio 16 camera.

## RESULTS

### Construction of *C. thermalis* mutants

To test the effects of using 8V Chls for growth under WL and FRL, the *bciB* gene of *C. thermalis* (Chro_1099) was disrupted with an erythromycin resistance cassette, generating a Δ*bciB* mutant (**Fig. 2A, B**). To act as a control for experiments with the Δ*bciB* mutant, as well as other mutants to be constructed in the future in *C. thermalis*, an additional mutant was generated in which the erythromycin resistance cassette was integrated into the genome between the converging Chro_1103 and Chro_1104 ORFs, ensuring that expression of these and neighbouring genes are not disrupted (**Fig. 2A, C**). This mutant, WT^Em^, could then be used for direct comparison with Δ*bciB* when selection with antibiotic was necessary.

**Figure 2.**
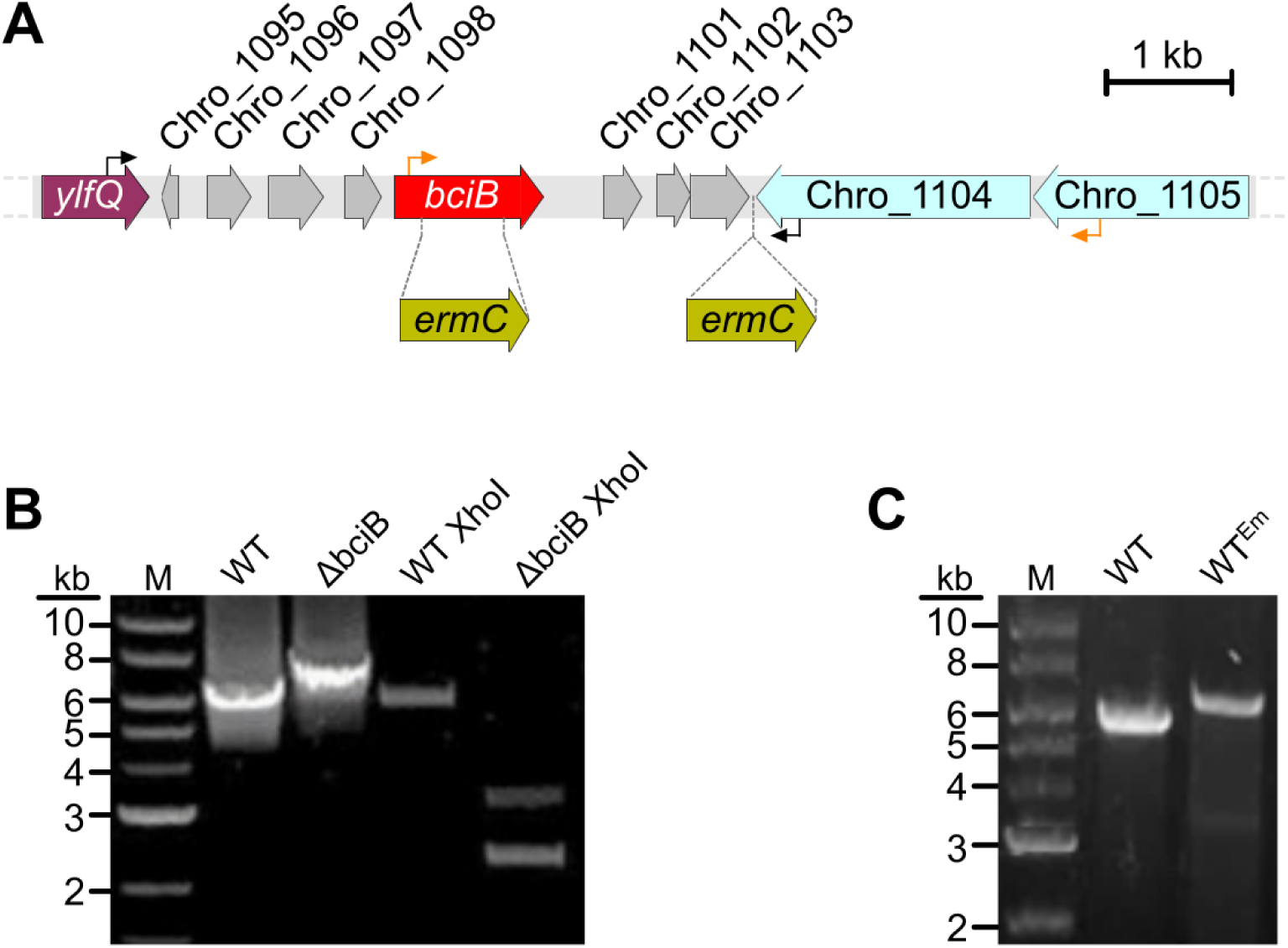
Construction of mutants of *C. thermalis*. *A*. Cartoon representation of the genomic location of the *bciB* gene. Sites targeted for the erythromycin resistance cassette (*ermC*) to disrupt *bciB* or generate a neutral site integration are indicated by dashed lines, and primer binding sites used to screen these mutations are indicated by black and orange arrows, respectively. *B*. PCR confirming the disruption of *bciB* from isolated genomic DNA. The WT amplicon does not contain a XhoI restriction site, so only that generated from the *bciB* mutant results in two shorter fragments of DNA after digestion with this restriction enzyme. *C*. PCR confirming the neutral site integration of the erythromycin resistance cassette, generating the WT^Em^ mutant, from isolated genomic DNA.

### Absorption profiles of *C. thermalis* strains

The WT^Em^ and Δ*bciB* strains of *C. thermalis* were cultured in liquid medium and illuminated by moderate intensity WL, and absorption spectra of whole cells were measured (**Fig. 3**, dashed-line spectra). When standardised by cell number, the Δ*bciB* mutant displays reduced amplitude peaks corresponding to photosynthetic pigment–protein complexes, indicating their decreased assembly in these cells. The Δ*bciB* mutant displays 8 nm and 1 nm red-shifts in Soret and Q_y_ absorption bands of the Chl-containing complexes, respectively, when compared to WT^Em^. When these strains were cultured moderate intensity FRL for 12 days, allowing sufficient time for them to have undergone the slow FaRLiP process, far-red absorbing features in the whole cell spectra were apparent (**Fig. 3**, solid-lined spectra, highlighted box). The absorption maxima displayed by the WT^Em^ strain were consistent between growth under WL and FRL, but growth under FRL results in an increase in the ratio of Chl-containing complexes to phycobilisomes in this strain. In comparison to WT^Em^, the Δ*bciB* mutant again displayed peaks associated with the photosystems and phycolbilisome antenna with reduced amplitude, including the far-red absorbing feature, and the mutant displayed a 7 nm red-shift in the Soret absorption band, but surprisingly also showed a Q_y_ band blue-shifted by 4 nm relative to WT^Em^, and by 5 nm compared to its maximum when grown under white light. The mutant also displays a decrease in the ratio of Chl-containing complexes to phycobilisomes when grown under FRL compared to WL, the opposite effect that growth under FRL elicits on WT^EM^.

**Figure 3.**
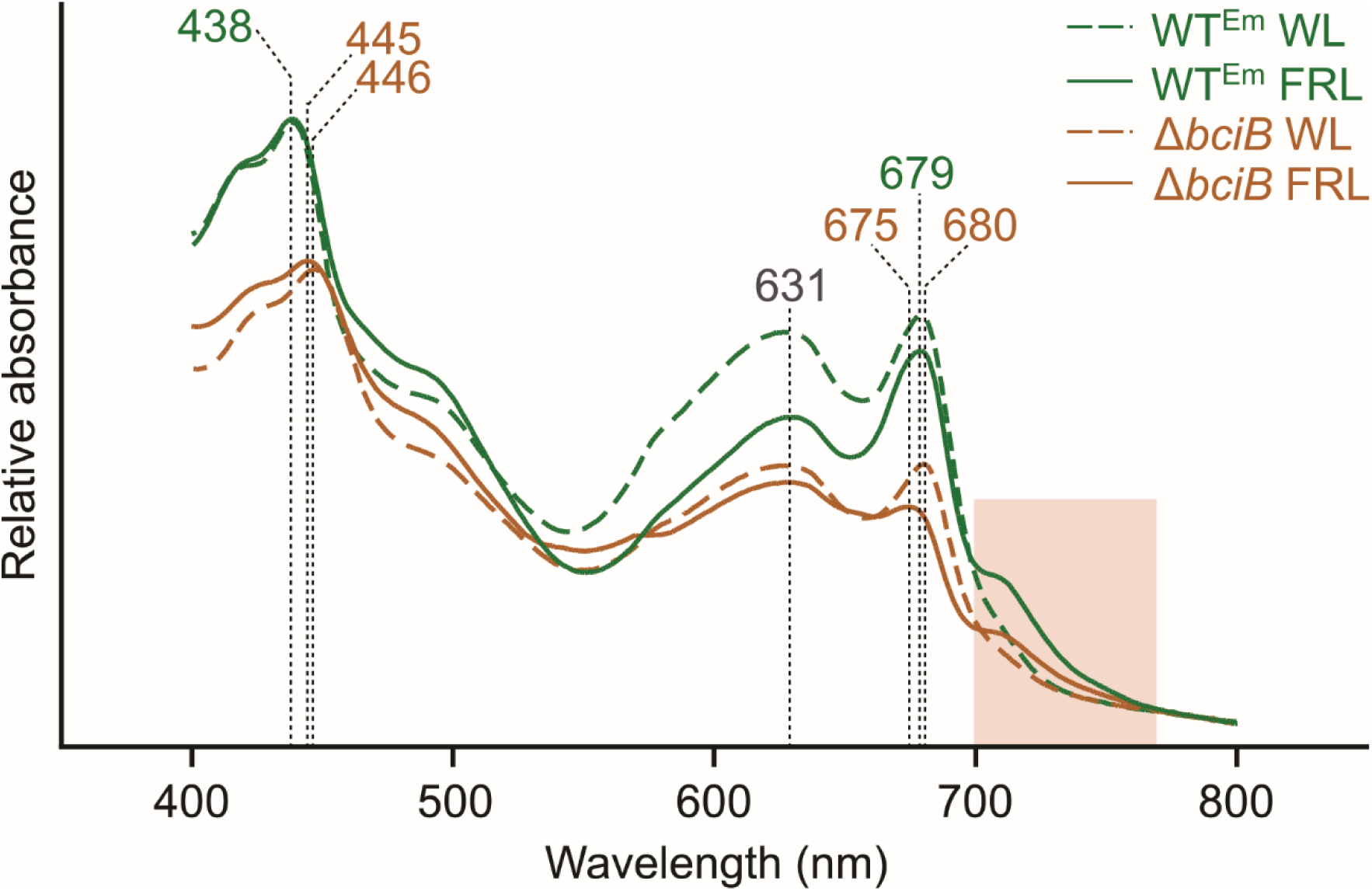
Whole cell absorption spectra of *C. thermalis* cells grown under WL and FRL. Cells from liquid cultures grown under moderate light were homogenised to break up clumps of cells, standardised by OD_780_, and their absorption spectra recorded. Absorption maxima of pigment-containing complexes are indicated by dashed lines, and the region from 700–770 nm in which spectral features appear in FaRLiP-capable cyanobacteria is highlighted.

### Pigment biosynthesis under white light and far-red light

Pigments were extracted from liquid cultures of WT^Em^ and Δ*bciB* grown under both WL and FRL and subjected to analysis by reverse-phase HPLC (**Fig. 4**). When grown under WL, WT^Em^ produces Chl *a* (carrying the canonical 8E group), while Δ*bciB* accumulates 8V-Chl *a*, identifiable by its shorter retention time and red-shifted Soret maximum in its absorption spectrum. Under FRL, WT^Em^ synthesises Chls *f* and *d* in addition to Chl *a*. The Δ*bciB* mutant also accumulates two additional pigments, each with a shorter retention time than their equivalent extracted from WT^Em^. The absorption spectra of these peaks demonstrate red-shifted Soret maxima, and in the case of the Chl *f* species, a blue-shifted Q_y_ maximum to 703 nm from 706 nm. These characteristics allow us to assign these pigments as 8V-Chls *d* and *f*, which have not been found in nature, to date. The spectrum of 8V-Chl *d* contains peaks at 439 and 665 nm, contributed from a minor Chl *a*-type species (perhaps esterified with a C15 tail); these pigments could not be separated to deconvolute their individual spectra despite employing a range of methods and columns. Chl *f* makes up ∼10% of the total Chls in WT^Em^ grown under FRL, consistent with other studies on FaRLiP cyanobacteria [32,33]. This contribution is reduced to ∼3% of total Chls in the Δ*bciB* mutant, indicating that the FaRLiP response is impaired when 8V-Chls are synthesised.

**Figure 4.**
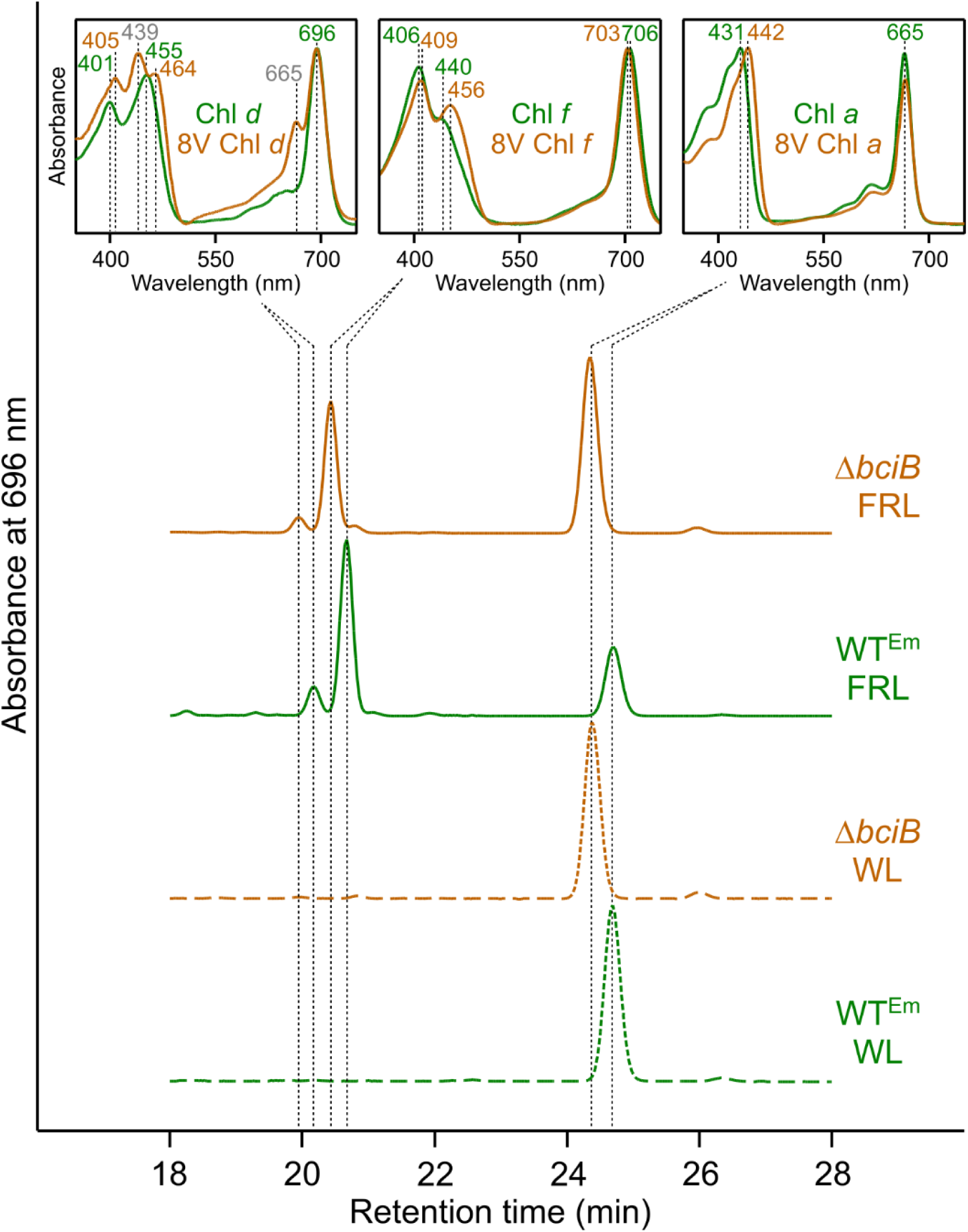
HPLC analysis of pigments extracted from described strains of *C. thermalis.* Pigments were extracted from liquid cultures grown in moderate intensity WL (dashed traces) and FRL (solid traces). In-line absorption spectra of peaks are shown in the inset panels, with absorption maxima used to identify the pigment species. Traces are normalised to major peak height for clarity.

### Energy transfer in whole cells

Whole-cell fluorescence emission spectra at 77 K were recorded for both strains (**Fig. 5**). When Chls were excited at 440 nm, WL grown cells display a major emission maximum at 726 nm and a broad, long-wavelength emission band at ∼800 nm, thought to derive from excitonically-coupled Chls at the monomer interface of dimeric and tetrameric PSI, which predominate in WL grown *C. thermalis* [34] (**Fig. 5A**). The Δ*bciB* strain displays prominent emission features with maxima at 641 nm, 666 nm, and 685 nm, the first two of these likely derived from phycocyanin and allophycocyanin, respectively, and the last from PSII. The features associated with allophycocyanin and PSII are detectable in the WT^Em^ spectrum, but are much less apparent. However, a conserved feature emitting at 695 nm, which has been associated with PSII [35], is more pronounced in WT^Em^ than in the Δ*bciB* cells.

**Figure 5.**
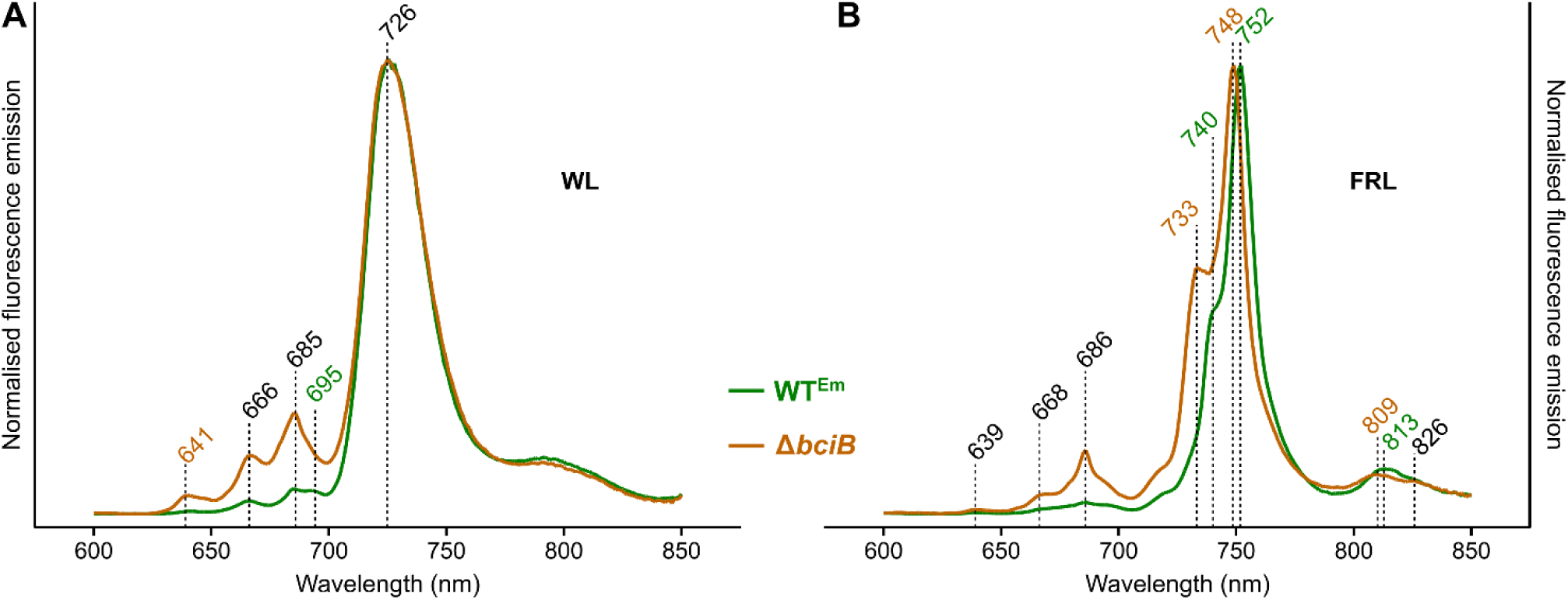
Low-temperature fluorescence emission spectra of *C. thermalis* cells grown under WL and FRL. Fluorescence emission spectra of WT^Em^ (green) and Δ*bciB* (orange) cells at 77 K, excited at 440 nm. **A**, Cells grown in WL. **B**, Cells grown in FRL. The emission intensities are normalised to those of PSI maxima, and emission features present in both strains, WT^Em^ only, and Δ*bciB* only are indicated in black, green, and orange, respectively.

When cells grown under FRL were excited at 440 nm, the major emission peak in WT^Em^ is shifted to 752 nm, derived from FRL-PSI, with a prominent shoulder associated with FRL-PSII at 740 nm (**Fig. 5B**). The equivalents in the Δ*bciB* mutant are blue-shifted by 4 nm and 7 nm, respectively. A similar blue-shift is observed for the PSI associated emission above 800 nm. The peak at 813 nm in WT^Em^ is shifted to 809 nm in the Δ*bciB* mutant. The emission band at 686 nm associated with WL-PSII is also still apparent in the Δ*bciB* mutant, which may be indicative of impaired and/or incomplete acclimation to FRL.

### Testing the light-sensitivity of *C. thermalis*

8VR mutants of *Synechocystis* and *A. thaliana* are sensitive to high-intensity WL [14,21]. Due to aseriate (clumping) morphology of *C. thermalis*, drop growth assays on solid medium were conducted to determine if the same effect is observed with this cyanobacterium, rather than monitoring absorbance of liquid cultures (**Fig. 6**). Under moderate light conditions, both WT^Em^ and Δ*bciB* perform equally well, indicating that production of 8V-Chl *a* does not impart sensitivity at this light intensity. However, at 250 μmol photons·m^−2^·s^−1^, Δ*bciB* cells are completely bleached, while the WT is able to grow, albeit inhibited when compared to the moderate light regime.

**Figure 6.**
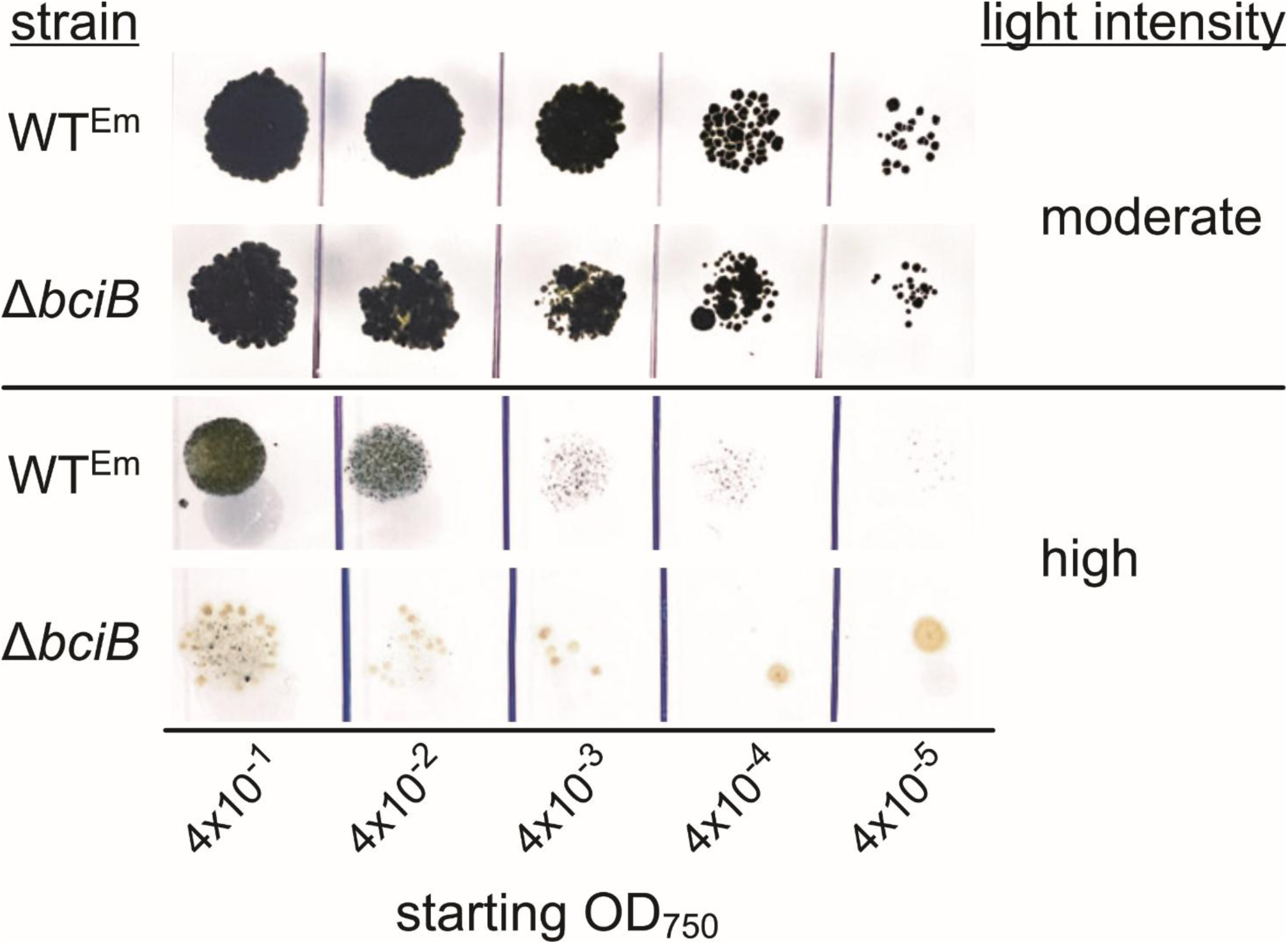
Drop growth assays of *C. thermalis* at moderate and high intensity WL. Clumps of cells from liquid cultures grown at moderate light were manually dispersed in a Dounce homogeniser, standardised by OD_750_ and serially diluted before spotting onto solidified BG-11 agar.

Light sensitivity under FRL was also tested. Since growth under this condition is too slow to be performed on plates, mid-exponential phase cells cultured under moderate WL were transferred to moderate and high FRL and allowed to undergo FaRLiP. Under moderate FRL conditions, those used to obtain the absorption spectra in Fig. 3, liquid cultures of WT^Em^ and Δ*bciB* are green, with no clear signs of bleaching (**Fig. 7**). However, when cultures are transferred to high intensity FRL to undergo FaRLiP, the Δ*bciB* cells bleach completely, and cannot be recovered when transferred back to moderate intensity WL (not shown). These results indicate that a similar sensitivity to high intensity illumination is displayed by *C. thermalis* cells under WL and FRL conditions when 8V Chls are synthesised.

**Figure 7.**
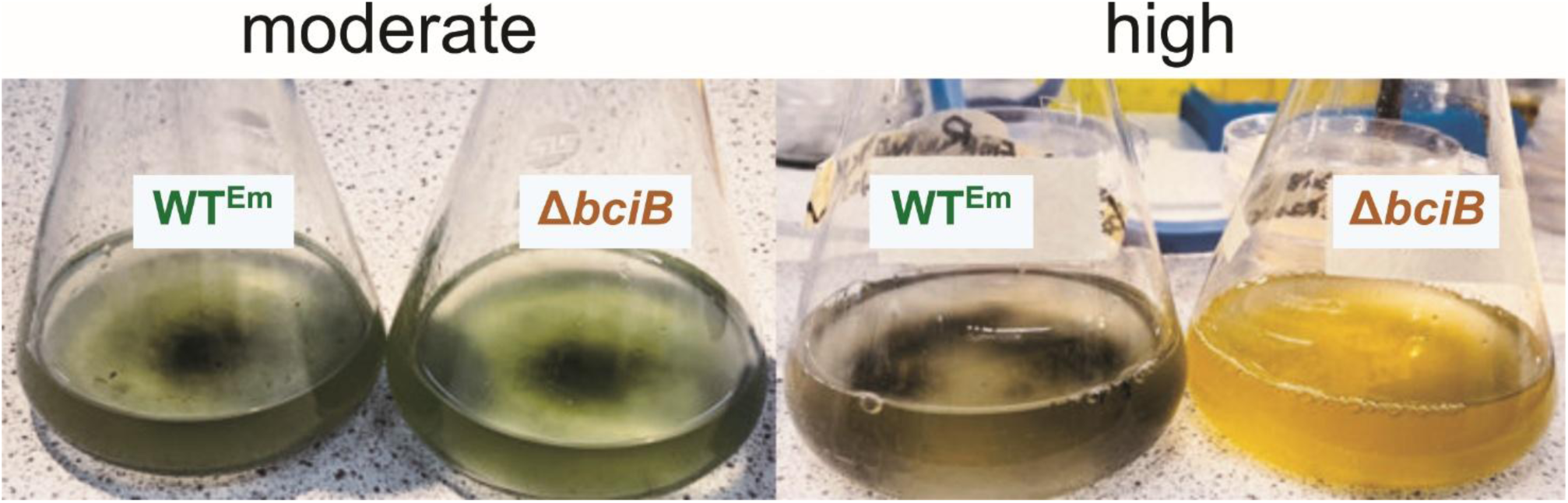
Liquid cultures of *C. thermalis* grown under moderate and high intensity FRL. Photographs were taken after 12 days of incubation under the relevant condition to allow the FaRLiP process to take place.

### Analysis of membrane ultrastructure

To determine if membrane morphology is affected by the synthesis of 8V-Chls, transmission electron micrographs of thin sections of *C. thermalis* cells were taken (**Fig. 8**). Cells were cultured under moderate intensity WL or FRL to mid exponential phase to provide moderate light samples. To generate samples exposed to high intensity light, additional mid exponential phase cells grown under moderate light were shifted to high intensity light for 2 days, allowing for bleaching effects to be observed while limiting cell death. WT^Em^ cells display a curved thylakoid membrane morphology under each condition, with overall cellular membrane density decreasing as light intensity increases under both WL and FRL conditions. This reduction in overall membrane density is also displayed by the Δ*bciB* mutant as light intensity increases, but this is accompanied by fragmentation of the thylakoids at high light, with uninterrupted membranes spanning the entire cell missing in these cells. The curvature of membranes in the Δ*bciB* mutant is also different to that of WT^Em^; cells synthesising 8E-Chls display thylakoids that bend into the cytoplasm, increasing overall surface area, while those in the mutant synthesising 8V-Chls remain contiguous with the cytoplasmic membrane.

**Figure 8.**
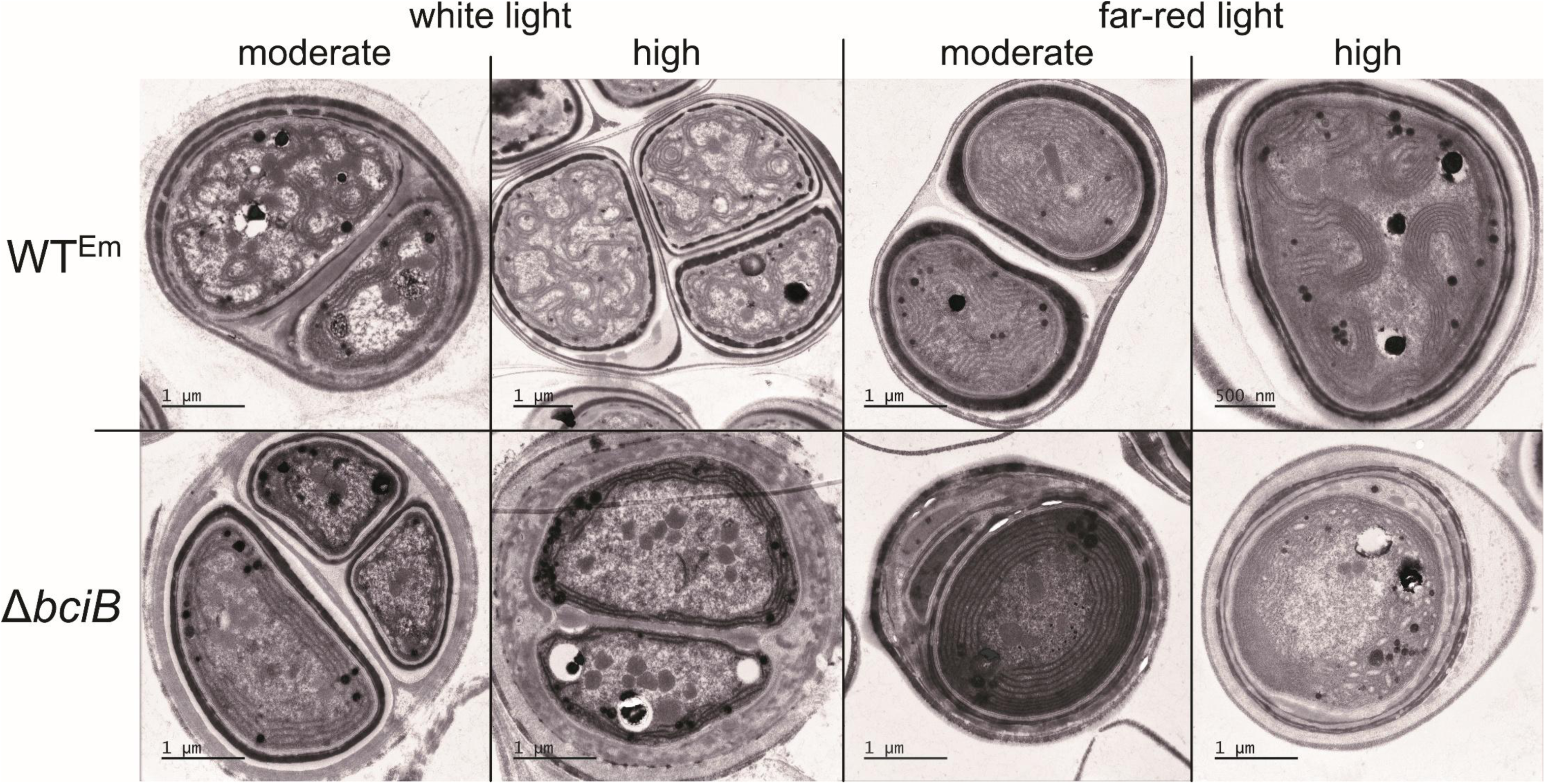
Transmission electron micrographs of thin sections of *C. thermalis* cells, grown under moderate and high WL and FRL. Electron microscopic images from WT^Em^ cells (upper panels) and Δ*bciB* cells (lower panels).

## DISCUSSION

In the present study we have determined that the synthesis of 8V-Chls results in light sensitivity under both WL and FRL in the model FaRLiP cyanobacterium *C. thermalis*. The results under WL mirror those reported for mutants of *Synechocystis* and *A. thaliana* accumulating 8V-Chls [21,14], but we report the first analysis of growth using 8V Chls *d* and *f* under FRL. Our results indicate that FaRLiP is impaired in the *bciB* mutant, with a much less prominent FRL-absorbing feature present in the whole-cell absorption spectrum, and a concomitant reduction in the Chl *f*:Chl *a* ratio, than is found in WT^Em^ cells.

Under both WL and FRL conditions, the *bciB* mutant accumulates fewer Chl-containing photosystems than WT^Em^, as indicated by the whole-cell absorption spectra. Similarly, under each condition tested, the *bciB* mutant displays an altered membrane morphology, where the characteristic bending thylakoids that curve into the centre of the cell, found in unmodified cells under both WL and FRL, are lost [34]. This lack of bending results in a reduction in the overall surface area of membrane in which to house PSII and PSI, and accounts for the loss in amplitude of the Chl-complex associated feature in the absorption spectra, but the direct cause of this change in morphology is not immediately obvious. The change in membrane shape is somewhat reminiscent of the effect of deletion of *curT* in *Synechocystis* [36]; the protein encoded by this gene is involved in thylakoid curvature in plants and cyanobacteria [37]. The genome of *C. thermalis* contains three orthologs of *curT*, which may in some part account for the extreme curvature of the membranes in this organism. However, further work is required to directly link synthesis of 8V-Chls to membrane morphology in the *bciB* mutant, including the fragmentation of these membranes under high-light conditions.

The high-intensity FRL employed in the current study may not be biologically relevant for certain environments where FaRLiP cyanobacteria are found (e.g. stromatolite, caves, microbial mats), where irradiance never reaches levels tested herein. However, high-intensity FRL may be experienced by cells growing under the surface layer of salt flats, where WL is almost completely reflected and FRL transmits, and under the canopy of plants, where FR wavelengths make up more than half of incident photons between 400–700 nm [38]. *Chroococcidiopsis* spp. are often found in hot desert environments, where photon flux can be extremely high [39]. It may be that FaRLiP cyanobacteria adapted to low-irradiance environments may better tolerate synthesis of 8V-Chls under these conditions.

It is notable that the clumping morphology of *C. thermalis* is less pronounced when cultured under FRL than under WL, which may permit a greater number of cells in a colony to access FRL. Under full spectrum sunlight, clumped colonies of FaRLiP-capable cyanobacteria may contain both cells performing canonical WL phototrophy and others that are limited to using FRL. It has been reported that *Chlorogloeopsis fritschii* PCC 6912, a FaRLiP cyanobacterium that also has a clumping morphology, produces Chls *d* and *f* when cultured in natural sunlight, but not when cultured under WL [40]. This is probably an effect of self-shading by the outer cells of the clump, and FRL being enriched as it penetrates the colony. If the common morphology of *C. thermalis* was retained under FRL, a similar scenario would occur, with the outer cells of clumps harvesting photons in the 700–780 nm range and restricting light availability to cells in the centre; a switch to a more planktonic morphology allows a greater number of cells to access FRL, but which may come at a cost of easier predation or loss of particulate organic carbon from the centre of the clump [41]. Interestingly, this is in contrast to the behaviour of *Acaryochloris marina*, which switches from a planktonic state when grown under WL to forming biofilms when cultured under FRL, although pigment biosynthesis in this organism is not altered during this switch [42].

*Prochlorococcus*, thought to be the most abundant genus on Earth, use 8V Chls *a* and *b*, naturally lacking 8VR enzymes. Many *Prochlorococcus* spp. are found in surface water, where photon flux can reach very high intensities [43]. It has been proposed that methionine 205 and cysteine 282 residues in the D1 (PsbA subunit of the PSII reaction centre (numbered according to *Thermosynechococcus vestitus* BP-1)— conserved in the *Prochlorococcus* D1, where valine and glycine are conserved, respectively, in 8E-Chl *a* cyanobacteria—play a role in the ability of these organisms to tolerate high light [44]. A *Synechocystis* mutant synthesising 8V-Chl *a*, modified to carry these *Prochlorococcus* residues gained light tolerance in an additive manner with V205M followed by G282C substitutions [44]. Interestingly, the C282 residue is found in the D1 subunit of Chl *d*-synthesising *Acaryochloris marina* strains, and is also found in the FRL-D1 protein produced when FaRLiP cyanobacteria are cultured under FRL (Figure S1). A structural alignment of the D1 subunits of WL-utilising *Synechocystis* [45] and of the FRL-PSII used by *Synechococcus* sp. PCC 7335 [46], along with Alphafold models of FRL-D1 from *C. thermalis* and D1 from the type strain of *Prochlorococccus* (*P. marinus* MIT 9313) [47] are shown in **Fig. 9**. The residues in these positions are in close proximity to the P_D1_–P_D2_ special pair and Pheo_D1_ pigments involved in photosynthetic electron transfer, but do not directly coordinate these cofactors. It could be that the bulkier methionine in place of valine at position 205 in the *Prochlorococcus* subunit reduces a steric clash between a less flexible 8V group and the protein backbone, or could play a role in limiting photodamage, as suggested by Ito and Tanaka [44]. A target of future work will be to make the V205M mutation in the FRL-D1-encoding *psbA3* gene in Δ*bciB* and WT^Em^ strains of *C. thermalis* to determine if enhanced tolerance to high intensity FRL can be conferred when using 8V-Chls, and what the penalties of the mutation are when using 8E-Chls, respectively.

**Figure 9.**
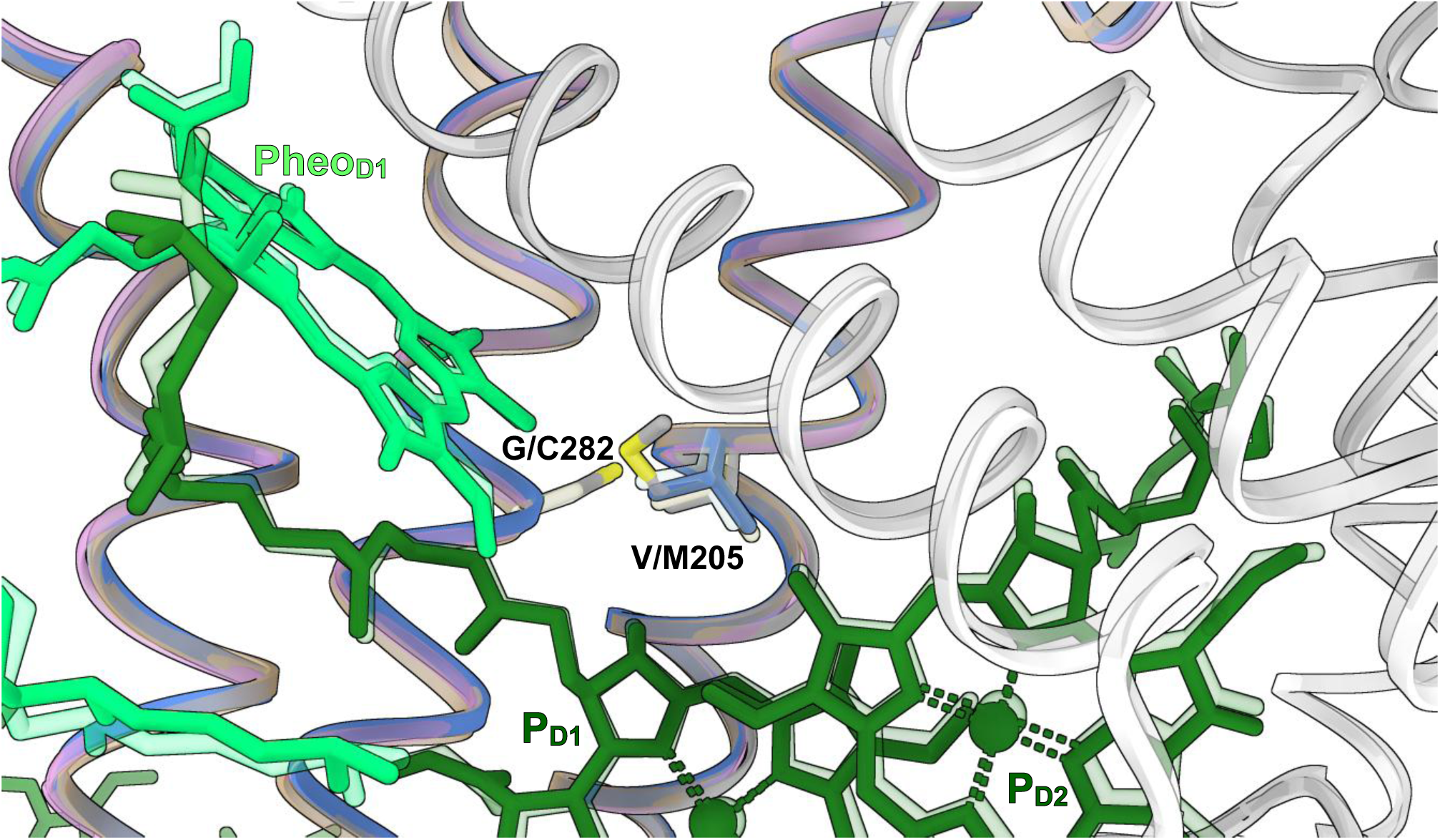
Structural alignment of D1 proteins. WL D1 of *Synechocystis* sp. PCC 6803 (7N8O; cornflower blue), FRL-D1 from *Synechococcus* sp. PCC 7335 (7SA3; tan), and the AlphaFold predictions of FRL-D1 from *C. thermalis* (K9TVY6; plum) and D1 from *P. marinus* MIT 9313 (Q7TTH6; silver). D2 proteins from *Synechocystis* sp. PCC 6803 (white) and *Synechococcus* sp. PCC 7335 (grey) are also shown. Chls (P_D1_ and P_D2_) and Pheo in the *Synechocystis* sp. PCC 6803 complex are shown in dark and light shades of green, respectively, and those from *Synechococcus* sp. PCC 7335 are shown at 80% transparency.

Our study has revealed that the synthesis of 8V-Chls affects growth, photosystem assembly, and membrane morphology under WL and FRL conditions in the FaRLiP cyanobacterium *C. thermalis*, and details the first biosynthesis of 8V Chls *d* and *f*. Despite 8V reduction being the only dispensable step in the biosynthesis of Chls in oxygenic photoautotrophs, the importance of 8VRs for FaRLiP and growth under high intensity light has been revealed. Future work will focus on characterising the photosynthetic performance of isolated 8V-Chl-containing photosystems assembled under FRL conditions.

## AUTHOR CONTRIBUTIONS

AKC, LKF, KP, and GFD data curation; AKC, KP, KP, GFD, DJN, RG and DPC formal analysis; DPC writing–original draft; AKC, LKF, KP, GFD, DJN, and RG writing–review and editing; RG and DPC supervision; DPC funding acquisition.

## FUNDING AND ADDITIONAL INFORMATION

This work was supported by grants from the Biotechnology & Biological Sciences Research Council, awards BB/W008076/1 (DPC and LKF) and BB/X015955/1 (DPC). AKC was supported by a University of Liverpool Studentship; DJN by the Deutsche Forschungsgemeinschaft (DFG; Emmy Noether project award no. NU 421/1). The authors gratefully acknowledge the University of Liverpool Biomedical EM Facility for their support and assistance in this work.

## DATA AVAILABILTY

The data that support the findings of this study, and materials used herein, are available from the corresponding author upon request.

