## Supplementary material for "Synthesis of C8-vinyl chlorophylls *d* and *f* impairs far-red light photoacclimation and growth under far-red light"

### SUPPLEMENTARY TABLES & FIGURES

| Strain/Plasmid | Properties | Source/Reference |
|---|---|---|
| *E. coli* | | |
| NEB® 5-alpha | Cloning strain for pRL277 constructs | New England Biolabs |
| HB101 | Donor strain for biparental mating | Promega |
| *C. thermalis* | | |
| WT | PCC 7203 | Pasteur Culture Collection |
| Δ*bciB* | Replacement of central ~700 bp of Chro_1099 with *ermC* from pRL692 in WT, $Em^R$ | This study |
| pseudoWT | Introduction of *ermC* from pRL692 into neutral site in WT genome, $Em^R$ | This study |
| Plasmid | | |
| pRL692 | Source of *ermC* cassette, $Sp^R$, $Em^R$ | Addgene, [1] |
| pRL277 | Cargo vector, $Sp^R$ | Addgene, [2] |
| pRL277[bciB KO] | pRL277 containing regions up- and downstream of Chro_1099 flanking *ermC*, $Sp^R$, $Em^R$ | This study |
| pRL277[pseudoWT] | pRL277 containing convergent regions of Chro_1103 and Chro_1104 flanking *ermC*, $Sp^R$, $Em^R$ | This study |
| pRL443 | Conjugal plasmid for mobilization of plasmids to cyanobacteria, $Tc^R$, $Ap^R$ | Addgene, [3] |
| pRL528 | Helper plasmid for bacterial conjugal DNA transfer, $Cm^R$ | Addgene, [4] |

**Supplementary Table 1. List of strains and plasmids described in this study**


[1] Koksharova OA, Wolk CP. 2002. A novel gene that bears a DnaJ motif influences cyanobacterial cell division. *Journal of Bacteriology*. **184**(19):5524-5528.

[2] Cai YP, Wolk CP. 1990. Use of a conditionally lethal gene in *Anabaena* sp. strain PCC 7120 to select for double recombinants and to entrap insertion sequences. *Journal of Bacteriology*. **172**(6):3138-3145.

[3] Elhai J, Vepritskiy A, Muro-Pastor AM, Flores E, Wolk CP. 1997. Reduction of conjugal transfer efficiency by three restriction activities of *Anabaena* sp. strain PCC 7120. *Journal of Bacteriology*. **179**(6):1998-2005.

[4] Elhai J, Wolk CP. 1988. Conjugal transfer of DNA to cyanobacteria. *In* Methods in enzymology (Vol. 167, pp. 747-754). Academic Press.

| Primer name | Sequence (5’-3’) | Restriction site |
|---|---|---|
| bciBKOUpF | GTGCGGCCGGTTAGCTATCTGCGATGATATTGG | EagI |
| bciBKOUpR | CAATTCTTTCAATGACTACACCCCTAGGCACCTCGAGCAATGGTACTAACAATACCCGTC | AvrII, XhoI |
| bciBKODownF | GACGGGTATTGTTAGTACCATTGCTCGAGGTGCCTAGGGGTGTAGTCATTGAAAGAATTG | XhoI, AvrII |
| bciBKODownR | GTCGAGCTCCTTATACAACTACTCTTGATAACC | SacI |
| pseudoWTUpF | GTGCGGCCGGACGGGTATTGTTAGTACCATTG | EagI |
| pseudoWTUpR | CTCTTTTGTTGCTTTGGCAGGCCTAGGCACCTCGAGGAGTGTTAGAAACTAGGCTG | AvrII, XhoI |
| pseudoWTDownF | CAGCCTAGTTTCTAACACTCCTCGAGGTGCCTAGGCCTGCCAAAGCAACAAAAGAG | XhoI, AvrII |
| pseudoWTDownR | GTCGAGCTCCTACAGAAGGAACCCTCATTACTG | SacI |
| EmRF | GTCACTCGAGGCGTGCTATAATTATACTAATTTTATAAGGAGG | XhoI |
| EmRR | CTGCCTAGGTGAGTGAGCTGATACCGCTCGCC | AvrII |
| bciBKOCheckF | GCACCCGGGGTAATTCCGAC | |
| bciBKOCheckR | CCTAAACTGGAAGCCGCGACC | |
| pseudoWTCheckF | GTGCATCAATCGATGACAGCGG | |
| pseudoWTCheckR | CTGAGCCAGCCAAGAACTATGG | |

**Supplementary Table 2. List of primers used in this study**

Restriction enzyme cleavage sites used for cloning are highlighted in the primer sequence.

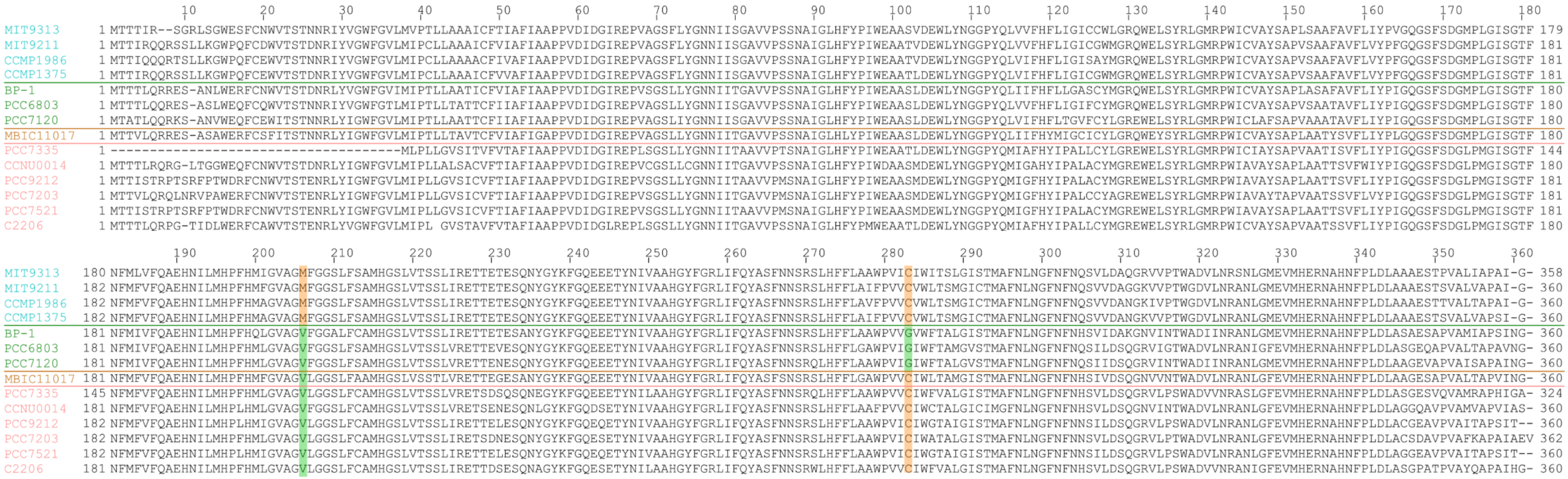


**Figure S1. Multiple sequence alignment of PSII D1 proteins, highlighting residues implicated in tolerance of 8V Chls.** D1 sequences from 8V-Chl producing *Prochlorococcus* spp. (cyan), model WL cyanobacteria that synthesise Chl *a* only (green), an *Acaryochloris* strain that primarily makes Chl *d* (orange), and FR PSII from FaRLiP cyanobacteria (pink), are shown, separated by horizontal lines. Culture collection/accession numbers are used in place of full names, and are ordered as follows: *Prochlorococcus marinus* MIT 9313, *Prochlorococcus marinus* MIT 9211, *Prochlorococcus marinus* subsp. pastoris CCMP 1986 (previously MED4), *Prochlorococcus marinus* subsp. pastoris CCMP 1986 (previously SS120), *Thermosynechococcus vestitus* BP-1 (formerly *T. elongautus* BP-1), *Synechocystis* sp. PCC 6803, *Nostoc* sp. PCC 7120 (aka *Anabaena* sp. DCC D0672), *Acaryochloris marina* MBIC11017, *Synechococcus* sp. PCC 7335, *Altericista leshanensis* CCNU0014, *Chlorogloeopsis fritschii* PCC 9212, *Chroococcidiopsis thermalis* PCC 7203, Fischerella thermalis PCC 7521 (aka *Mastigocladus laminosus* Y-16-m), *Halomicronema hongdechloris* C2206.

,
